# Restricted proliferation during neurogenesis contributes to regionalization of the amphioxus nervous system

**DOI:** 10.1101/2021.12.22.473870

**Authors:** Giacomo Gattoni, Toby GR Andrews, Elia Benito-Gutiérrez

## Abstract

The central nervous system of the cephalochordate amphioxus consists of a dorsal neural tube with an anterior brain. Two decades of gene expression analyses in developing amphioxus embryos have shown that despite the lack of overt segmentation the amphioxus neural tube is highly regionalized at the molecular level. However, little is known about the mechanisms that generate such precise regionalization. Proliferation is a key driver of pattern formation and cell type diversification, but in amphioxus it has never been studied in detail nor in the specific context of neurogenesis. Here, we describe the dynamics of cell division during the formation of the central nervous system in amphioxus embryos and its contributions to the regionalization of the neural axis. We show that after gastrulation, proliferation pauses to become spatially restricted to the anterior and posterior ends of the neural tube at neurula stages. Only at the onset of larval life, proliferation resumes in the central part of the nervous system. By marking specific populations and inhibiting cell division during neurulation, we demonstrate that proliferation in the anterior cerebral vesicle is required to establish the full cell type repertoire of the frontal eye complex and the putative hypothalamic region of the amphioxus brain, while posterior proliferating progenitors, which were found here to derive from the dorsal lip of the blastopore, contribute to elongate the caudal floor plate. Between these proliferative domains, we find trunk nervous system differentiation is independent from cell division, which decreases during neurulation and resumes at the early larval stage. Taken together, our results highlight multiple roles for proliferation in shaping the amphioxus nervous system.

## Introduction

The central nervous system (CNS) of amphioxus has been the subject of intense study. This is due to the key phylogenetic position of amphioxus (cephalochordate) as the sister group to all other chordates, which makes them unique to investigate the evolutionary origin of vertebrate nervous systems (Garcia-Fernàndez and Benito-Gutiérrez, 2009; Benito-Gutiérrez, 2011; Holland, 2016; Holland and Holland, 2021). These studies have primarily employed two distinct approaches: first, through exquisitely detailed morphological descriptions of the late amphioxus larva and adult brain anatomy by electron microscopy (Lacalli et al., 1994; Lacalli, 1996; Wicht and Lacalli, 2005); and second, through molecular characterization of cell types, via analysis of gene expression patterns and protein localization in the CNS during embryogenesis and in adult amphioxus (Benito-Gutiérrez, 2006; Yu et al., 2007; Albuixech-Crespo et al., 2017; Holland, 2017). It is only recently however, that both approaches are being considered in an integrated way to understand the development and organisation of the amphioxus CNS (Benito-Gutiérrez et al., 2021; Lacalli, 2021). As a whole, such studies have revealed a high degree of molecular regionalisation within the amphioxus CNS, despite its relatively simple morphology. The amphioxus CNS consists of a hollow dorsal neural tube that, unlike vertebrate nervous systems, is not overtly segmented, with an anterior enlargement representing the “cerebral vesicle”(Wicht and Lacalli, 2005). Despite this simplicity, gene expression analysis has revealed numerous cell types which, albeit in low numbers, collectively represent regions of homology to specific areas of the vertebrate brain and spinal cord (Albuixech-Crespo et al., 2017; Benito-Gutiérrez et al., 2021).

During embryonic development, the emergence of complex cell type diversity occurs concomitantly with the physical processes of tissue morphogenesis. This means that cells differentiate in the context of tissues that are undergoing dynamic transitions in their shape, size and cellular architecture, by virtue of diverse processes including cell division, rearrangement and growth. Cell division exerts critical functions in tissue morphogenesis. This includes regulation of tissue-specific cell number and size (Weigmann et al., 1997; Lanet et al., 2013; Winkley et al., 2019), the orientation and magnitude of tissue growth and shape change (Baena-López et al., 2005; Pickering et al., 2018; Iber, 2021), and the emergence of cell type diversity through asymmetric distribution of cytoplasmic determinants (Cowan and Hyman, 2004; Williams and Fuchs, 2013; Almonacid et al., 2014). As such, the dynamics of cell division is a prime candidate for regulating the emergence of morphological complexity and diversity in animal evolution. An understanding of how modulation of cell division dynamics may have contributed to evolutionary transitions depends on characterisation of its properties in a wide diversity of taxa.

In vertebrates, proliferation has essential roles in multiple aspects of nervous system development, including neurulation, differentiation and post-embryonic growth (Qian et al., 2000; Wullimann and Knipp, 2000; Alvarez-Buylla and Garcia-Verdugo, 2002; Caviness et al., 2003; Sadler, 2005; Kaslin et al., 2008). A proper balance between proliferation and programmed cell death is key to establish final neuronal numbers and adult brain size (Rakic, 1995; Chenn and Walsh, 2002; Ando et al., 2019). Furthermore, the total number of neurons seems to correlate with behavioural complexity and higher cognitive capabilities (Herculano-Houzel, 2017; Logan et al., 2018). In vertebrate embryos neuroepithelial progenitors and radial glial cells proliferate in the ventricular zone of the neural tube. These progenitors can either divide symmetrically to augment the progenitor pool or asymmetrically to produce neuroblasts that differentiate into different types of neurons (Caviness et al., 1995; Götz and Huttner, 2005; Huttner and Kosodo, 2005). Although the balance and rate of symmetric versus asymmetric divisions is specific to different vertebrate species (Rakic, 1995; Kriegstein et al., 2006; Fish et al., 2008), in general, cells proliferate extensively across the entire CNS, explaining the high number of cells and the resulting larger size of vertebrate nervous systems (Frederikson and McKay, 1988; Caviness et al., 1995; Wullimann and Knipp, 2000).

In addition to its role in increasing neural complexity, cell division is required to define form and function in the vertebrate spinal cord. In early development, this is particularly obvious in the tailbud, where posterior axial progenitors expand clonally to contribute with new cells to the elongating neural tube (Wilson et al., 2009; Steventon and Martinez Arias, 2017). Axial progenitor cells arise from regions within, and adjacent to, the organiser at late gastrula stages, and ultimately relocate to the tailbud during axial elongation, in a region of contact between the posterior neural tube and axial mesoderm termed the chordoneural hinge (Selleck and Stern, 1991; Catala et al., 1996; Cambray and Wilson, 2002). This includes posterior midline progenitors, which contribute to elongation of the floor plate and notochord (Catala et al., 1996; Gray and Dale, 2010; Row et al., 2016) and neuromesodermal progenitors (NMps) which elongate the dorsolateral neural tube and paraxial mesoderm (Cambray and Wilson, 2002; Tzouanacou et al., 2009; Mugele et al., 2018). While progenitor cells with similar topology and fate have been identified in a diversity of vertebrate species, their clonal dynamics, and therefore their contributions to axial elongation, are highly variable (Steventon and Martinez Arias, 2017). For example, fate mapping studies in mouse and chick embryos have shown axial progenitors to make broad contributions to axial development, with progeny populating the entire post-occipital body axis (Cambray and Wilson, 2002; Albors et al., 2018; Mugele et al., 2018). In contrast, similar approaches in zebrafish have identified very little proliferation of axial progenitors within the tailbud, and contributions to only to the extreme tip of the tail (Attardi et al., 2019). This variation raises an important question of whether similar progenitor cell types exist in amphioxus, and if so, what contribution they make to formation and maturation of the spinal cord.

In amphioxus, previous studies have reported restricted proliferation in the anterior tip of the neural tube and the posterior floor plate at the mid-neurula stage (Holland and Holland, 2006). However, the origin, dynamics and fate of these proliferating cells, and therefore their contributions to the emergence of neural complexity in amphioxus, remain to be investigated. In this work, we present a detailed characterisation of cell division dynamics during early neural development in amphioxus. In doing so, we identify some critical roles of proliferation in the diversification of the neural type repertoire and in the specification of neural axis geometry. We first expand on previous data (Holland and Holland, 2006; Andrews et al., 2021) by constructing a quantitative proliferation landscape during early neural development, which shows that cell division restricts after gastrulation to distinct mitotic niches in the cerebral vesicle and chordoneural hinge. By arresting cell division pharmacologically, we identify unique roles for cell division in each territory. In the anterior cerebral vesicle, cell division is required for generating specific neuronal types and therefore increasing cell type diversity within the maturing brain. In the posterior neural tube, cell division is necessary to elongate the posterior floor plate and notochord, in a population of axial progenitors arising from the dorsal blastopore lip of the gastrula. Collectively, our data shows that cell division has several critical roles during the formation of the nervous system in amphioxus, thereby suggesting that fine modulation of cell division dynamics might have been a major strategy for the emergence of neural complexity through chordate evolution.

## Materials & Methods

### Animal husbandry and embryo fixation

Adults of the European amphioxus *B. lanceolatum* were collected in Banyuls-sur-Mer, France, and transported to Cambridge, UK, where they were maintained and spawned in a custom-made facility as described in (Benito-Gutiérrez et al., 2013). After *in vitro* fertilization, embryos were raised in filtered artificial salt water (ASW) at 21°C and fixed in ice-cold 3.7% Paraformaldehyde (PFA) + 3-(N-morpholino)propanesulfonic acid (MOPS) buffer for 12 hours, then washed in sodium phosphate buffer saline (NPBS), dehydrated and stored in 100% methanol (MeOH) at -20°C.

### EdU labelling and detection

For EdU pulse analyses, EdU was applied to live embryos in filtered seawater at a final concentration of 20μM for 2 hours prior to fixation. For EdU pulse-chase analyses, EdU was removed after 2 hours of exposure *in vivo* by transferring embryos to a fine 15μm filter and washing them in an excess of filtered sea water. They were then transferred to a fresh petri dish and incubated until the desired developmental stage.

Fluorescent detection of incorporated EdU was performed following the manufacturer’s instructions, using a Click-it EdU Alexa Fluor 647 Imaging Kit (Invitrogen) prior to primary antibody incubation. As advised for enhanced signal, the copper reagent was replenished after 15 minutes.

### Pharmacological perturbation with hydroxyurea

Live amphioxus embryos were treated with 2μM hydroxyurea (Sigma, H8627) or an equal volume of dimethylsulfoxide (DMSO; Sigma, 276855). This was performed either between the cup-shaped gastrula and 14-somite stages (8hpf to 34hpf at 21°C), or the 6-somite and the 12- and 14-somite stages (18hpf to either 30 or 34hpf at 21°C).

### Immunohistochemistry

Rehydrated embryos were permeabilised overnight in PBS + 1% DMSO + 1% Triton and incubated in a bleaching solution of 3% H2O2 + 3% formamide in 0.2X SSC. Embryos were then blocked in PBS + 0.1% Triton + 0.1% BSA + 5% NGS for 3 hours. The blocking solution was then replaced, including primary antibodies as follows: rabbit anti-laminin (Sigma, L9393) at 1:50, rabbit anti-PhH3 (Abcam, ab5176) at 1:500, mouse anti-acetylated tubulin (Sigma, T6793) at 1:250. Primary antibody incubation was performed overnight at 4°C, followed by washes in PBS + 0.1% Triton + 0.1% BSA and then by a secondary block of PBS + 0.1% Triton + 0.1% BSA + 5% NGS for 3 hours. Finally, this was replenished, to also include goat anti-rabbit and/or goat anti-mouse secondary antibodies at 1:250 and DAPI at 1:500 for overnight incubations. Embryos were washed thoroughly with PBS + 0.1% Triton. Imaging was performed on an Olympus V3000 inverted confocal microscope.

### *In situ* hybridization chain reaction (HCR) on embryos

HCR version 3 was performed on embryos as described in (Andrews et al., 2020). Briefly, embryos were rehydrated in NPBS + 0.1% Triton X, incubated for 30 minutes in bleaching solution and permeabilized in 1% DMSO, 1% Triton for 3 hours. They were incubated in Hybridization Buffer (HB, Molecular Instruments) for 2 hours and then probes were added in HB overnight at 37°C. The following day probe excess was removed with Wash Buffer (Molecular Instrument). Embryos were washed in 5X-SSC + 0.1% Triton X, incubated in Amplification Buffer (AB, Molecular Instruments) for 30 minutes and then left overnight in the dark at room temperature in AB + 0.03μM of each hairpin (Molecular Instruments). The next day embryos were washed in the dark in 5X-SSC + 0.1% Triton X and incubated overnight with 1 μg/mL DAPI in NPBT, then washed in NPBT and transferred in a glass-bottomed dish in 100% glycerol. Imaging was performed on an Olympus V3000 inverted confocal microscope.

### Image analysis

To overcome EdU signal saturation from the endoderm, nuclear EdU labelling was masked using a binary mask of the DAPI channel in Fiji. Neurons and axonal projections were manually segmented using the Surface tool of the Imaris software (IMARIS 9.7.2, Bitplane, Oxford Instruments).

## Results

### Cell division restricts to two polarized domains during neural tube morphogenesis

To define the pattern of cell division during amphioxus neural tube formation, we first constructed a spatiotemporal map of cell cycle progression between gastrulation (0-somite stage; 2 hours prior to formation of the first somite) and the early larva stage (14-somite stage) (Figure 1A). At each somite-stage across this time course, we labelled embryos with markers for nuclei in two cell cycle phases. We incubated embryos in EdU for 2-hours prior to fixation, detecting cells passing through S-phase, and immunostained for phosphorylated histone 3 (PhH3), to mark mitotic nuclei at the time of fixation. To synthesise data across multiple specimens, we calculated the mean frequency of labelled nuclei in evenly sized bins of the anteroposterior axis at each stage, and plotted these values across stages and mean axial lengths to generate a proliferation ‘landscape’. This map of cell cycle progression revealed cell division in the neural tube to pass through multiple phases, exhibiting unique spatial dynamics (Figure 1A). First, at the end of gastrulation, it becomes specific to the extreme posterior neural plate, where it interfaces with the posterior axial mesoderm at the dorsal blastopore lip (Figure 1A 0 – 4 ss, 1B 0ss + 3ss). Second, between 5 ss and 10 ss, we observed a burst of cell division restricted to the extreme anterior and posterior tips of the neural tube (Figure 1A 5 – 10 ss, 1B 7 ss + 10 ss). At these stages, we identified almost no cell division in the central part of the neural tube, between these polarised mitotic domains. Finally, after 10 ss, cell division resumed at a low level across the entire anteroposterior axis (Figure 1A 10 ss – 14 ss, 1B 12 ss + 14 ss).

**Figure 1.**
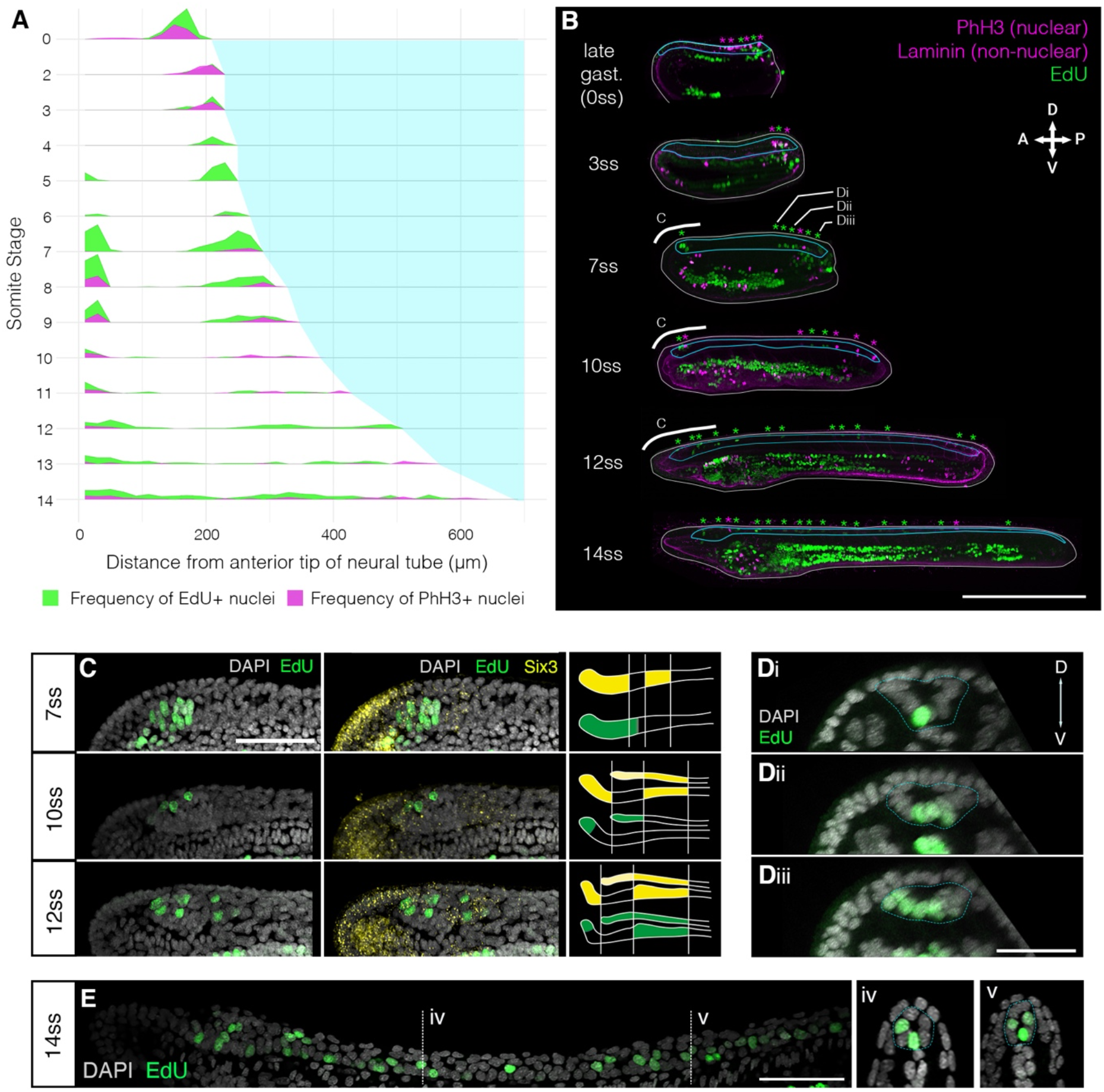
Cell division restricts to two polarised mitotic domains during neural tube morphogenesis. **(A)** Proliferation landscape for the neural tube, showing mean frequency of EdU-positive (EdU exposure for 2-hours before fixation) and PhH3-positive nuclei in 20μm bins of the anteroposterior axis at successive somite stages. ‘0-somite stage’ refers to late gastrula 2-hours before onset of somitogenesis. n = 114 embryos. **(B)** Mid-sagittal sections for representative embryos included in (A), showing distribution of EdU-positive and PhH3-positive nuclei in each tissue of the body axis, which are delineated by laminin immunostaining. White lines outline the whole embryo, light blue lines outline the neural tube. Asterisks mark location of EdU/PhH3-positive nuclei using the same color code. Scale bar is 150μm. **(C)** Lateral z-projections of the cerebral vesicle at 7, 10 and 12 somite stages, at the level indicated by thick white lines in B, showing EdU incorporation and co-localisation with *Six3/6*, using HCR in situ hybridisation. Scale bar is 50μm. Brain patterns are schematically represented and summarised on the right side of the panel in the same color code **(D)** Serial transverse sections through a 7-somite stage embryo at the posterior positions illustrated in B (Di, Dii, Diii), showing the EdU detection profile. Neural tube is outlined with a dashed blue line. Scale bar is 30μm. **(E)** Lateral projection of 14-somite stage embryo, with representative transverse sections at the indicated positions: iv and v, where the neural tube is outlined with a light blue dashed line. Scale bar is 50μm.

Having identified a spatial restriction of cell division to the anterior and posterior poles of the neural tube, we next sought to define the pattern of cell division in each domain in finer detail. For the anterior domain, we combined EdU detection with HCR for *Six3/6*, a known marker of anterior neuroectoderm (Steinmetz et al., 2010; Range, 2014). During neurulation, *Six3/6* expression resolves the amphioxus cerebral vesicle into three areas: anterior and posterior *Six3/6*-positive domains, separated by a small region devoid of *Six3/6* expression, what we term here as intercalated *Six3/6* negative region (Kozmik et al., 2007; Albuixech-Crespo et al., 2017). Our analysis showed that each of these areas proliferates at different rates during neural tube morphogenesis. At 7 ss, EdU labelled a relatively large group of cells in the anterior *Six3/6*-positive and intercalated Six3/6 negative region, whereas no EdU-positive cells were found in the posterior *Six3/6-*positive cluster (Fig1C). At 10 ss, cell division declined, such that only 3-5 EdU-positive cells were identified in the anterior tip or dorsal region of the developing cerebral vesicle, within the anterior *Six3/6*-positive cluster (Figure 1C). At 12 ss, the number of EdU-positive cells in the anterior-dorsal cerebral vesicle increased, and proliferating cells became also visible in the caudal cerebral vesicle within the posterior *Six3/6*-positive domain (Figure 1C).

To characterize the spatial distribution of cell division in the posterior body, we analysed the pattern of EdU-positive nuclei in serial transverse sections from whole mount images along the anteroposterior axis. This was performed at 7 ss, when cell division is occurring at its greatest magnitude in the posterior neural tube, and EdU-positive are seen extending anteriorly from the chordoneural hinge (Figure 1D). At 7 ss, the posterior neural plate has not yet folded to form a tube and exhibits a ‘U’ shape in transverse section. The most anterior EdU-positive cells in the neural tube were invariably located ventrally, in the prospective floor plate (Figure 1D i). However, in more posterior sections, EdU-positive cells were more dispersed across the mediolateral axis of the neural plate, such that multiple EdU-positive cells were observed in each transverse section (Figure 1D ii). In sections immediately anterior to the chordoneural hinge, EdU-positive cells were found throughout the neural plate, with no conspicuous spatial bias across the mediolateral axis (Figure 1D iii). Considered together, cell division occurs broadly in the posterior neural plate during axial elongation, but restricts to the most medial-ventral cells at more anterior axial positions.

At 14 ss, when cell division has resumed in the area between the two proliferative domains, we found that EdU labelled cells were located only in the floor plate or in medio-lateral cells, clearly visible in cross-sections of the neural tube (Figure 1E). In sum, this analysis reveals that cell division restricts during axial elongation to two polarised mitotic domains; the anterior cerebral vesicle, where different regions of the prospective brain exhibit different proliferative rates, and the posterior neural tube, where proliferative cells extend anteriorly from the chordoneural hinge into the posterior floor plate.

### Proliferation specifically contributes to maturation of the brain and posterior floor plate

We next sought to map the contributions of proliferative cells in the anterior and posterior mitotic domains of the neural tube to tissue morphogenesis. We employed a pulse-chase approach to mark proliferative cells in each domain when the polarised cell division dynamic is conspicuous (5 ss – 10 ss) and map the distribution of their clones at 12 ss when the polarised proliferation dynamic has diminished. Pulses of EdU at 7-8 ss labelled distinct clusters of cells at the anterior tip of the neural tube (Figure 2Ai), and a single-file row of cells in the posterior neural tube extending anteriorly from the chordoneural hinge of the tailbud (Figure 2Aii). The ventral endoderm was also strongly labelled but was non-overlapping with the two neural proliferative domains. After the EdU pulse, EdU was washed out extensively with fresh sea water, and embryos were left to develop until the 12 ss. At 12 ss, we found that EdU-positive cells in the anterior remained confined to the cerebral vesicle, and did not spread posteriorly into the rest of the neural tube (Figure 2Bi). Meanwhile, EdU-positive cells in the posterior neural tube contributed to 1/3^rd^ of the floor plate (Figure 2Bii). Furthermore, some EdU-positive cells initially located in the chordoneural hinge at 7-8ss, were found in the posterior notochord at 12 ss (Figure 2Bii compare with Figure 2Aii).

**Figure 2.**
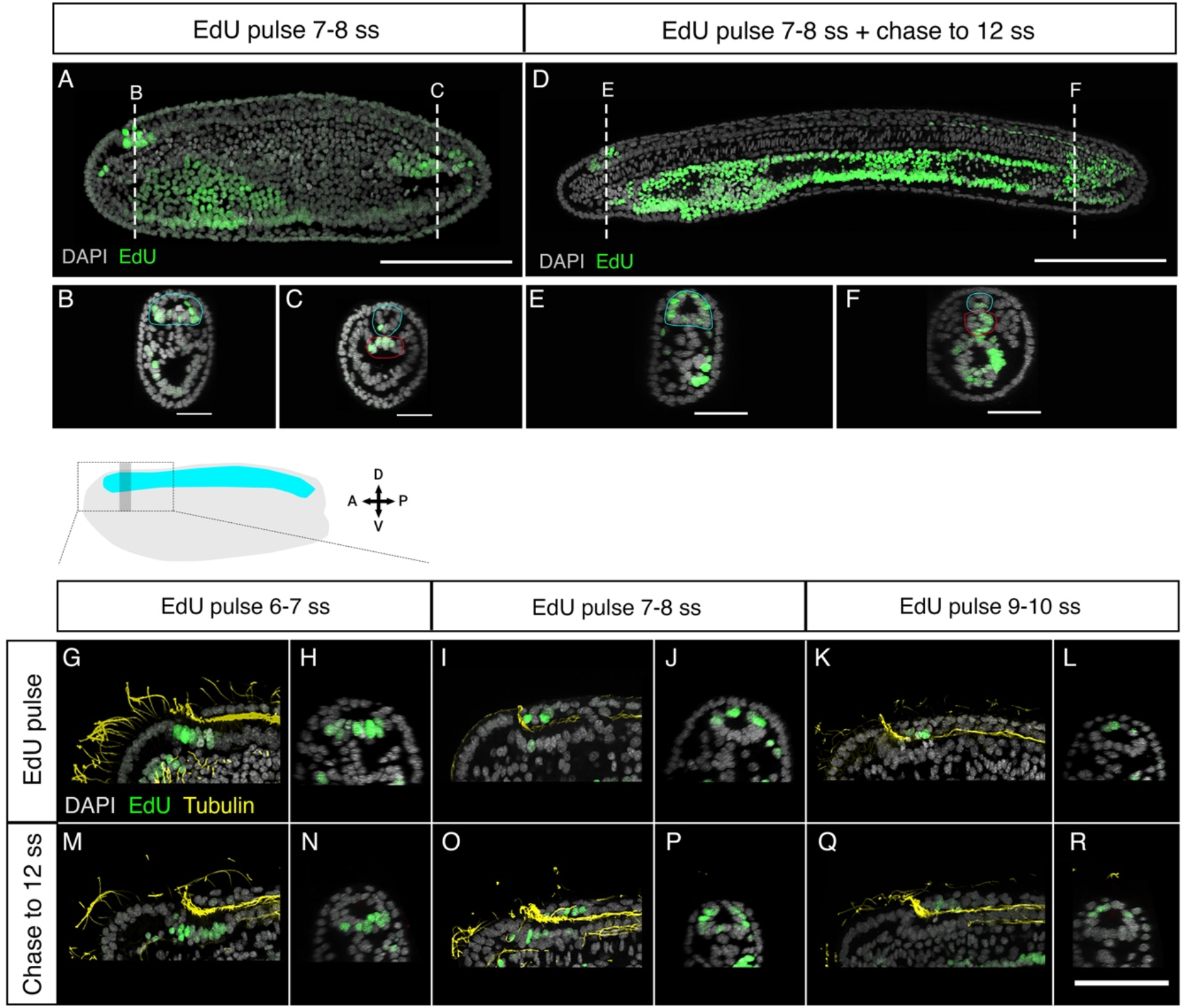
Fate of proliferating cells tracked using EdU-pulse chases. **(A)** Detection of EdU-pulse between 7-8 ss with cross section details shown for the anterior **(B)** and the posterior **(C)** neural proliferative domains. **(D)** Chase of the same EdU-pulse shown in A to 12ss revealing that while anterior EdU-positive cells remain in the brain **(E)** posterior positive cells spread through one-third of the posterior floor plate **(F)**. Scale bar is 100μm for lateral views and 30μm for cross-sections. **(G-H)** EdU pulses at 6-7ss marked ventral brain cells, as shown in lateral (G) and sagittal (H) planes, which remained ventral in the cerebral vesicle of 12ss embryos, as shown in lateral **(M)** and sagittal **(N)** planes of the brain. **(I-L)** At later stages lateral and dorsal proliferating cells in the brain, as shown by Edu pulses between 7-8ss (I: lateral plane; J sagittal plane) and 9-10ss (K: lateral plane; L sagittal plane) contributed to progressively dorsal cell populations, as shown in embryos at 12ss in lateral **(O, Q)** and sagittal **(P, R)** planes. EdU in green, acetylated tubulin in yellow. Scale bar is 50μm.

As we showed above, neural progenitors within the cerebral vesicle proliferate at different rates between 7 ss and 12 ss. We therefore performed pulse-chase experiments across this time window to determine how cell division contributes to the maturation of specific brain areas during development. Clones of cells that are in S-phase and incorporate EdU between 6 and 7 ss remained ventral at 12 ss, meaning they contribute broadly to the ventral side of the cerebral vesicle along its anteroposterior axis (Figure 2Ci). In contrast, EdU-positive cells labelled between 7 ss and 8 ss contributed most significantly to the lateral walls of the cerebral vesicle (Figure 2Cii), and those labelled between 9 ss and 10 ss contributed exclusively to the dorsal side of the cerebral vesicle (Figure 2Ciii). Considered together, these data suggest that cell division in the brain is spatially restricted, increasing cell numbers locally in a ventral-to-dorsal temporal sequence. Meanwhile, proliferative cells in the posterior neural tube and chordoneural hinge specifically contribute to elongation of the posterior floor plate and become broadly dispersed across the anteroposterior axis within the posterior body.

### Cell division is dispensable for broad body plan patterning but necessary for proper axial tissue geometry

After tracing the distribution and fates of proliferative cells in the amphioxus neural tube, we sought to assess the functional contributions of cell division to neural tube morphogenesis. To this end, we first inhibited cell division from gastrulation (8 hpf), when cell division is widespread throughout the embryo (Holland and Holland, 2006), to the early larva stage (14 ss) using hydroxyurea (HU) to arrest DNA synthesis (Figure 3A).

**Figure 3.**
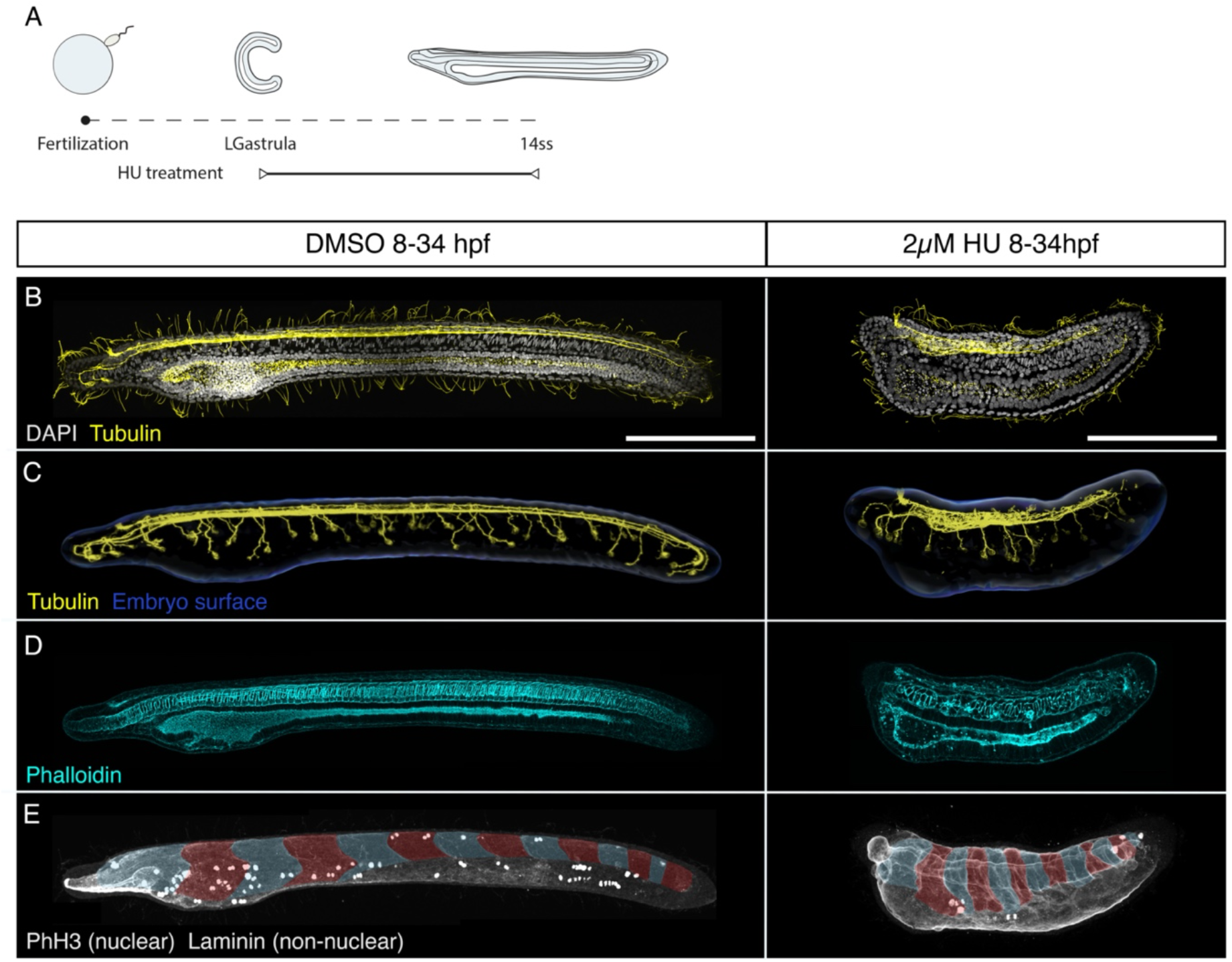
Cell division is dispensable for body plan patterning but required for establishing a normal geometry. **(A)** Experimental design of hydroxyurea (HU) treatments. **(B-E)** Control embryos (treated with DMSO) and embryos treated with 2μM HU are shown side by side in lateral z-projections. **(B)** The general morphology of these embryos is shown through immunostaining with acetylated tubulin (cilia and axonal scaffold) and DAPI (nuclei). **(C)** The structure of the axonal scaffold is shown by segmentation of the acetilated tubulin channel from embryos in (A). **(D)** Phalloidin (actin) staining in the same embryos reveals the presence of a notochord and a ventral endoderm. **(E)** Immunostaining with laminin and PhH3 reveals the presence of an equal number of somites in both control and treated embryos (false-colouredin blue and read overlay), and the efficacy of the treatment to arrest proliferation, which results in a significant loss of PHH3-positive nuclei in HU-treated embryos. Scale bars are150μm for both control and treated embryos. HU-treated phenotype representative of n = 13 embryos imaged.

Surprisingly, HU-treated embryos remained viable until the 14ss stage, hatching and swimming normally (data not shown) despite the lack of cell division (Figure 3). While cell death was widespread in HU-treated embryos, as indicated by pyknotic DAPI-stained nuclei (data not shown), the major patterns of the body axis were intact. The neural tube was present and internalized beneath the surface ectoderm (Figure 3B-C). Immunostaining for acetylated-tubulin in both control and treated embryos showed a central neural canal and axonal tracts running on the ventral side of the neural tube (Figure 3B-C). The neural tube of HU-treated embryos showed a clearly defined cerebral vesicle, which opened anteriorly through the neuropore and had rostral axonal projections similar to DMSO-treated control embryos. Furthermore, peripheral epidermal sensory neurons formed normally along the body axis and projected to the neural tube, although the size of each neuron was visibly larger than in control embryos (Figure 3C). These observations demonstrate the developmental robustness of the nervous system in amphioxus, which in essence forms and regionalises normally even in the absence of proliferation during early neurogenesis (Figure 3C).

Aside from the nervous system, the notochord was present at the axial midline in HU-treated embryos, with a characteristic stack-of-coins pattern, and a complete pattern of somites was formed within the paraxial mesoderm on the left and right sides of the embryo (Figure 3D-E). While this means that the amphioxus body plan is robust to the loss of cell division, our results also show that the geometry of the embryos is severely distorted. Most conspicuously, HU-treated embryos failed to elongate, reaching less than half the AP length of DMSO-treated siblings by 14 ss. Our results thus indicate that proliferation after gastrulation may be dispensable for the broad patterning of the body plan, but it is key to confer axial tissues with proper geometry.

### Anterior proliferation is required for cell type diversity of the brain

Having found that cell division is dispensable for formation of the neural tube and its broad regionalisation into cerebral vesicle and spinal cord, we next investigated its functions within each proliferative domain.

First, we focussed on the brain. Despite its morphological simplicity, the amphioxus brain is characterized by a complex diversity of cell types (Benito-Gutiérrez, 2006; Albuixech-Crespo et al., 2017). To test whether cell division is required to increase cell type diversity in the brain as opposed to only increment cell numbers, we treated embryos with HU starting from the 6ss stage, when proliferation is localised in the brain and chordoneural hinge, and raised these embryos to 12 ss and 14 ss (Figure 4A, Supplementary Figure 1A). We then examined the expression of genes known to mark specific cell types in chordate brains: the serotonergic marker *SerT*, the glutamatergic marker *VGlut* and the transcription factors *Six3/6* and *Otp* (Figure 4-5, Supplementary Figure 1B-D).

**Figure 4:**
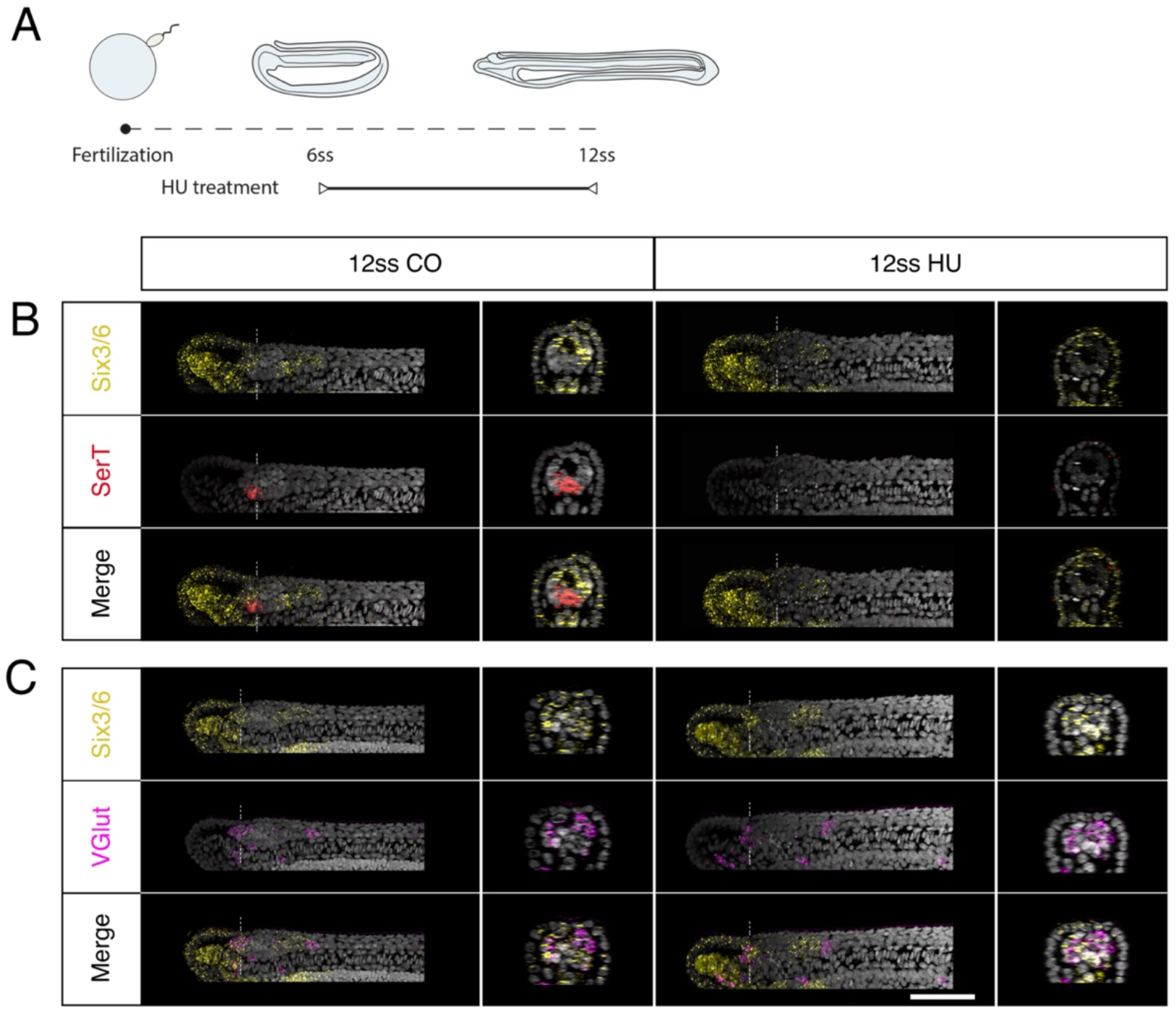
Brain proliferation is required to establish the neuronal type repertoire in the amphioxus larva. Expression of neurotransmitter markers in control and Hydroxyurea (HU)-treated embryos at 12 ss through HCR. **(A)** Experimental design of HU treatments. **(B)** *SerT* expression (red) in control embryos is localized at the border of the anterior *Six3/6* domain (yellow) but disappears following HU treatment. **(C)** In the neural tube, *VGlut* (magenta) is expressed in anterior-most cells within the anterior *Six3/6* domain and caudal to the posterior *Six3/6* domain. This pattern remains the same in HU-treated embryos. Scalebar is 50μm.

We observed no change in the tripartite organisation of the *Six3/6* expression domain (previously reported by Kozmik et al., 2007) in HU-treated embryos (Figure 4B-C). This observation was consistent with these domains forming prior to axial elongation and before the anterior cerebral vesicle starts proliferating, at stages before the HU was administered. By contrast, the small cluster of serotonergic (*SerT-*positive) neurons, characteristic of Row2 in the cerebral vesicle of 12ss control embryos (Candiani et al., 2012; Vopalensky et al., 2012), is absent from HU-treated embryos, suggesting that localised proliferation is needed to generate Row2 serotonergic neurons (Figure 4A, Supplementary Figure 1B). In control embryos, the glutamate transporter *VGlut* was expressed in the anterior *Six3/6*-positive domain at 12 and 14ss and caudal to the posterior *Six3/6*-positive cluster, as described previously (Candiani et al., 2012). Unlike *SerT*, the expression pattern of *VGlut* was not significantly altered by HU treatment (Figure 4B, Supplementary Figure 1C).

*Otp* is expressed in the hypothalamus, diencephalon and hindbrain in vertebrate embryos, where it contributes to specify a variety of neurons, including dopaminergic and peptidergic cells (Del Giacco et al., 2008; Fernandes et al., 2013). Our results confirm that in amphioxus *Otp* is expressed in the trunk region of the nervous system at the 6ss stage, as previously reported (Albuixech-Crespo et al., 2017) (Figure 5Ai). In addition, we further characterize *Otp* expression at later stages and find that *Otp* is expressed in the brain for the first time at 10 ss, in a pair of medial cells located in the posterior *Six3/6*-positive region (Figure 5Aii). At 12 ss new *Otp*-positive cells form two prominent ventral clusters in the intercalated *Six3/6*-negative region (Figure 5Aiii, B, C). Co-detection of Otp with Edu in 12 ss embryos pulsed at 7-8 ss, when ventral brain cells proliferate (See Fig2), confirmed these anterior Otp-positive clusters to be proliferating at 7-8ss (Figure 5H). In accordance with this result, while the pair of *Otp*-positive neurons in the posterior *Six3/6*-positive domain was also detected in HU-treated embryos at 12 ss, the clusters of *Otp*-positive neurons in the intercalated *Six3/6*-negative region were lost (Figure 5I). This supports a requirement for cell division in the specification of Otp cells at this particular location in the brain. In control embryos at 14 ss the number of *Otp* neurons in the posterior *Six3/6*-positive region increased (Figure 5Aiv). However, we found no co-localization of EdU and *Otp* at this specific stage in the brain or trunk region (Figure 5D-G). Prolonging the treatment with HU to 14 ss impeded the increase of *Otp-*positive neurons in the posterior Six3/6-positive domain that would be expected in normal conditions (Supplementary Figure 1D).

**Figure 5:**
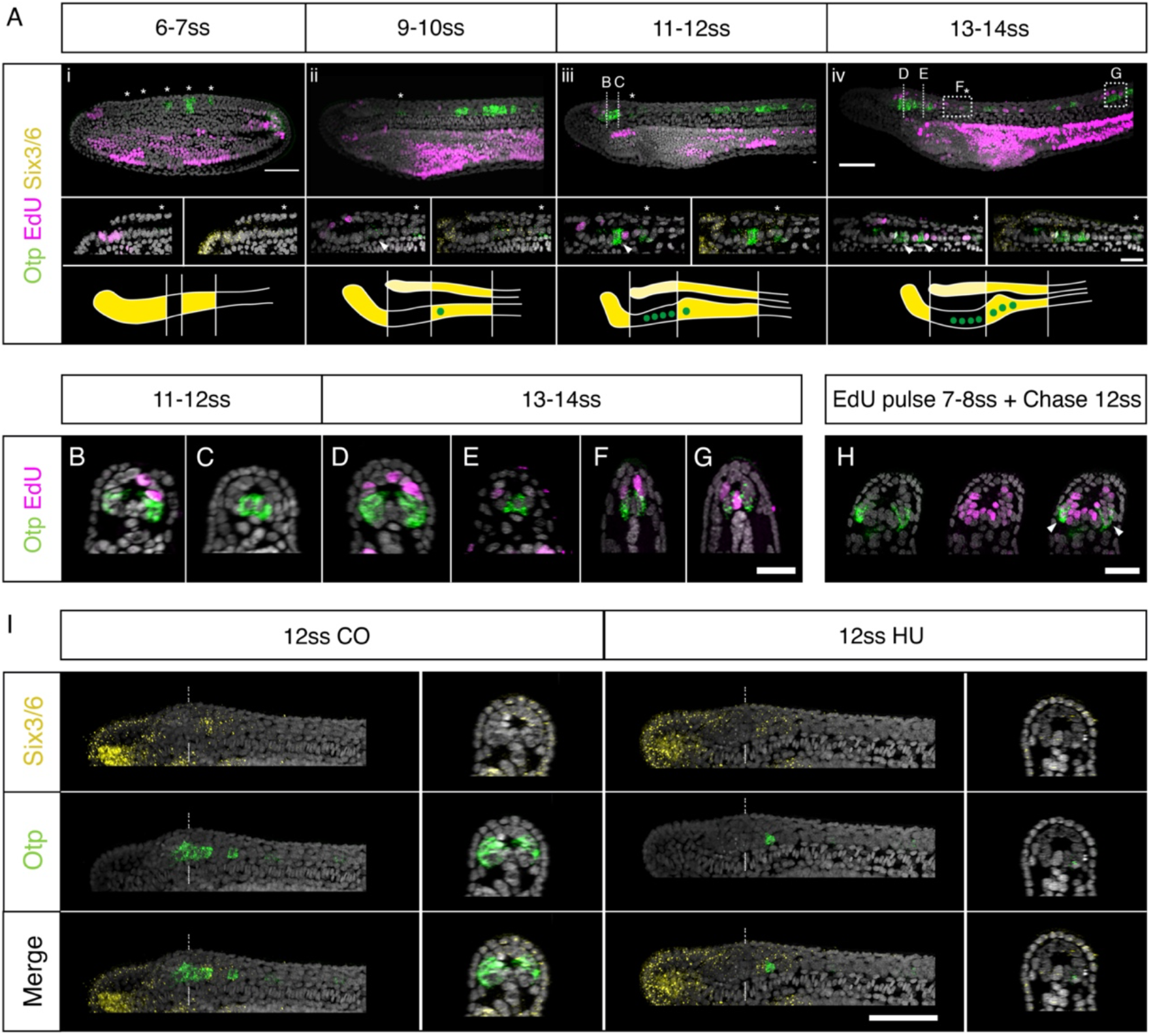
*Otp* expression and brain proliferation during neurulation. **(A)** Co-detection of *Otp* and *Six3/6* expression through HCR in situ hybridisation and EdU incorporation. **(i)** At 7ss *Otp* is expressed in 5 clusters of cells in the trunk region (asterisks). Expression is strong in the posterior three clusters but very low in the anterior two clusters (asterisk in insets). **(ii)** At 10 ss, a pair of *Otp* positive neurons appear in the posterior *Six3/*6 domain of cerebral vesicle (arrowhead in insets), but proliferation is still restricted to the anterior cerebral vesicle. **(iii)** At 12 ss two ventro-lateral clusters of *Otp*-positive cells are visible in the anterior cerebral vesicle. At this stage, cell division starts in the posterior cerebral vesicle; the posterior *Otp*-positive pair is not proliferating but cells adjacent to them are labelled by EdU (insets). **(iv)** At 14 ss, the number of posterior *Otp*-positive neurons increases (arrowhead). Asterisks in all insets show the first *Otp* cluster in the trunk. Patterns of Six3/6 and Otp co-expression are schematically representated at the bottom of each panel for every developmental stage. Scale bar is 50μm; 20μm for insets. **(B-G)** Cross section of 12 ss and 14 ss embryos showing the anterior ventro-lateral **(B**,**D)**, posterior medial **(C**,**E)** and trunk **(F**,**G)** *Otp* cells. Scalebar: 20μm **(H)** 12 ss embryos EdU-pulsed at 7ss show co-localization of EdU and *Otp* (arrowheads) at the level of D in Aiv. **(I)** Inhibition of cell division with 2μM hydroxyurea leads to a loss of the ventro-lateral *Otp* clusters in the *Six3/6*-negative domain, while the posterior medial *Otp* cells are not affected. Scale bar is 50μm.

Considered together, these data indicate that cell division timely regulates the specification of particular brain cell types at distinct time points, thereby incrementing the neural cell type diversity throughout development.

### Posterior midline progenitors have a common origin in the dorsal blastopore lip

Having identified a role for cell division in generating cell type diversity in the brain, we next focused on its contribution in the posterior neural tube, where we found cell division expands the floor plate. We have previously shown that proliferation starting at the 6 ss is required for full elongation of the body axis (Andrews et al., 2021). However, little is known about the progenitor cells that fuel axis elongation in amphioxus. Fate mapping studies in vertebrates have revealed axial progenitor cells residing in the chordoneural hinge of the tailbud to derive from the dorsal blastopore lip of the late gastrula, or its equivalent in amniotes, the node-streak border. During axial elongation, a subset of these axial progenitors clonally expands, and specifically contributes new cells to the posterior notochord and floor plate. We therefore sought to ask if a similar progenitor population exists in amphioxus.

Since direct cell marking and tracking *in vivo* is technically challenging in amphioxus, we further took advantage of the highly localised proliferation dynamics we identified here to spatially map the origin of chordoneural hinge progenitors in the gastrula, prior to axial elongation. Closer observation of EdU incorporation in the blastopore of the late gastrula revealed a temporal delay in cell cycle progression across the dorsoventral axis. In EdU pulses between 10 hpf and 12 hpf, EdU was primarily incorporated by cells in the dorsal and upper-lateral lips of the blastopore (Figure 6A-C, G). In contrast, EdU pulses between 12 hpf and 14 hpf marked cells in the ventral and lower-lateral lips of the blastopore (Figure 6D-F, J). Based on this observation, we reasoned that we could determine the derivatives of each blastopore subdomain by performing successive pulse-chase experiments, following the sequential enrichment of EdU across the dorsoventral axis of the blastopore.

**Figure 6.**
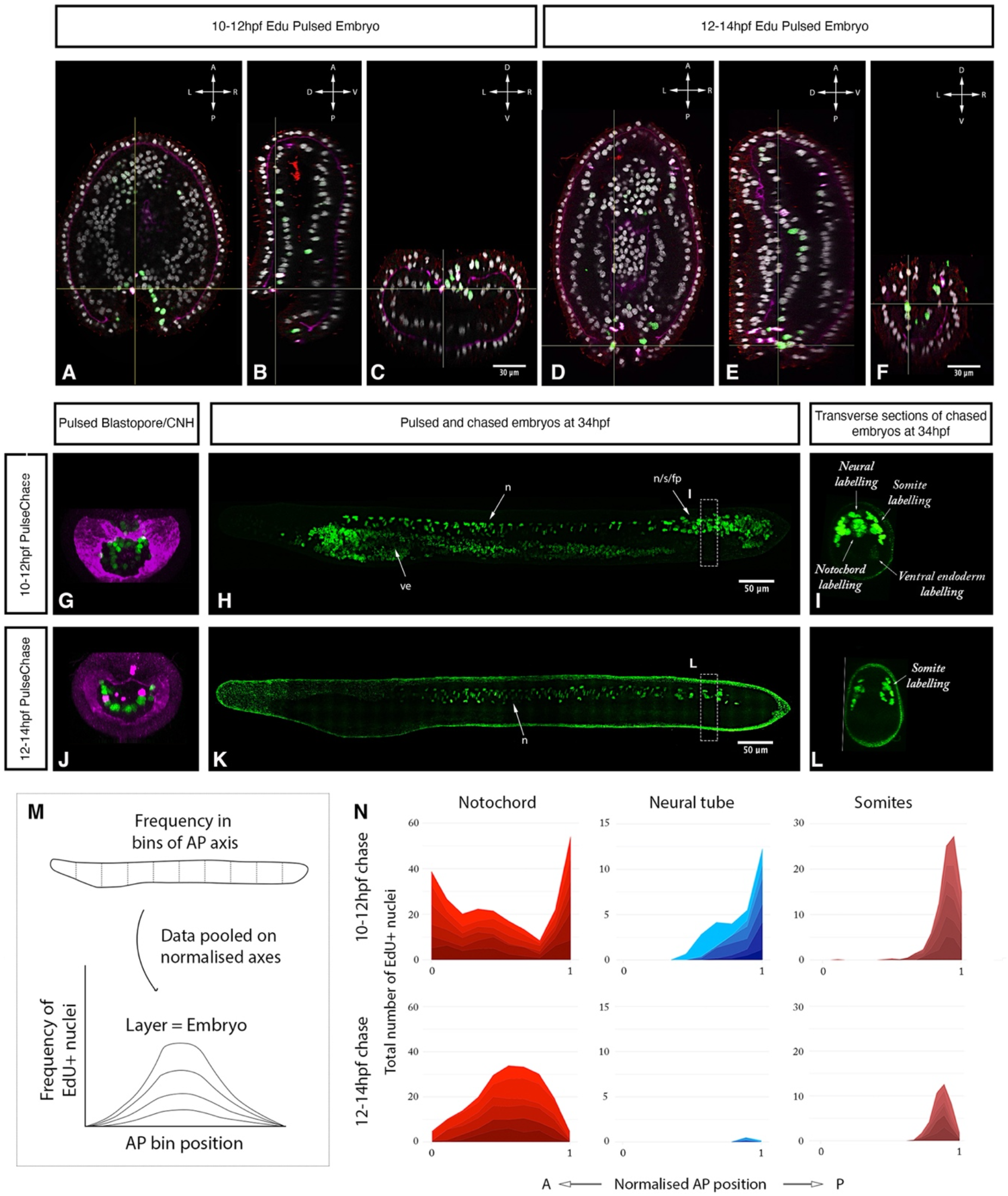
Axial progenitor fates are spatially regionalised across the late blastopore lip. **(A-C)** Distribution of EdU- and PhH3-positive nuclei after an EdU pulse between 10hpf and 12hpf, shown in orthogonal coronal (A), sagittal (B) and transverse sections (C) intersecting at the level of the blastopore (as indicated by the yellow lines). **(D-F)** Distribution of EdU-positive nuclei after an EdU pulse between 12hpf and 14hpf, shown in a coronal (D), sagittal (E) and transverse sections (F) intersecting at the level of the blastopore (as indicated by the yellow lines). Compass shows orientation of each plane in (A-F). **(G)** Maximum projection through the blastopore in an embryo pulsed with EdU between 10hpf and 12 hpf, as shown in (C), marking proliferative cells in the dorsal and upper-lateral lips. **(H)** Maximum projection through the sagittal midline of an embryo EdU pulsed at 10-12hpf, as shown in (G), showing contribution of EdU-positive cells in the dorsal blastopore lip to neural tube, notochord and somites. **(I)** Transverse section through posterior body of embryo shown in H, showing tissue-specificity of EdU-positive cells. **(J)** Maximum projection through the blastopore in an embryo pulsed with EdU 12-14 hpf, as shown in F, showing proliferating cells in the ventral and lower-lateral lips. **(K)** Maximum projection through the sagittal midline of an embryo EdU pulsed at 12-14hpf, as shown in J, reavealing the contribution of proliferating cells to the posterior somites. **(L)** Transverse section through posterior body of embryo shown in K, showing tissue-specificity of EdU-positive cells. **(M)** Schematic of approach used to quantify EdU labelling in pulse-chased specimens as a frequency curve across normalised AP length, following quantification in 10 evenlysized bins. **(N)** Stacked area graphs for each pulse-chase condition and tissue. n = 6 embryos per condition. EdU in green, PhH3 (nuclear) and laminin in magenta, acetylated tubulin in red. Scale bars as indicated. Abreviattions: fp, floor plate; s, somites; n, notochord; ve, ventral endoderm.

In the first of these pulse-chase experiments (EdU pulse 10 – 12 hpf), we found that cells initially located in the dorsal and upper-lateral blastopore lips at 12 hpf made broad contributions to the posterior body when chased to 14 ss (Figure 6H). At 14 ss, EdU-positive cells were enriched in the chordoneural hinge of the tailbud, and extended anteriorly from it to populate the posterior floor plate, notochord and posterior 4-5 somites (Figure 6H, I). This is shown qualitatively in midline sections of representative 14 ss embryos (Figure 6H), and quantitatively by calculating the frequency of EdU-positive nuclei in evenly-sized bins of the anteroposterior axis in each axial tissue (Figure 6M, N). Labelling of the chordoneural hinge, floor plate and notochord in this experiment recapitulated the distribution we noted previously in a pulse-chase experiment with EdU incorporation at 7-8 ss when the chordoneural hinge is proliferating (Figure 2C). This finding therefore suggested that progenitor cells of the chordoneural hinge have an origin in the blastopore lip of the late gastrula.

In the second pulse-chase experiment (EdU pulse 12 – 14 hpf), we found that cells initially located in the ventral and lower-lateral blastopore lips of the late gastrula (Figure. 6J) gave rise to posterior somites at 14 ss, with almost no EdU-positive cells detected in the posterior notochord or floor plate (Figure 6K, L). In this second pulse-chase experiment, loss of EdU labelling in the dorsal blastopore lip during the EdU pulse (compare figure 6G and 6J) correlated with a loss of labelling in the posterior notochord and floor plate at 14 ss (compare figure 6I, L). By taking advantage of temporal changes in blastopore proliferation dynamics, this experiment therefore indicated a spatial regionalisation of progenitor subtypes in the late blastopore lip, which includes: a) a dorsal lip containing progenitors that relocate to the chordoneural hinge of the tailbud and enter a proliferative phase after 6 ss to elongate the posterior floor plate and notochord; and b) lateral and ventral lips containing progenitors that give rise to cells populating the posterior 4-5 somites.

## Discussion

Proliferation is a key regulator of morphological complexity in nervous system development, with elucidated functions in the regulation of tissue size, geometry, and cellular complexity. As such, the number of cell divisions in neural progenitors and the balance between proliferation and differentiation exhibits striking variation between species (Kriegstein et al., 2006; Fish et al., 2008; Montgomery, 2017; Logan et al., 2018; Briscoe and Ragsdale, 2019). While vertebrate species are characterized by large brains, including anatomically distinct compartments and complex stratification of neural cell types, the two groups of invertebrate chordates - cephalochordates and tunicates – possess simpler central nervous systems, composed of fewer cells and exhibiting limited morphological regionalization (Sasakura et al., 2012; Holland and Holland, 2021). A key question is what role cell division plays in the morphogenesis of these simple nervous systems, and whether these dynamics may predict the emergence of more complex properties in vertebrates.

### Landscapes of cell division in amphioxus show developmental modularity during the formation of the CNS

In this study, we provide new insight into the distribution and function of cell division in the embryonic nervous system of amphioxus. We expand on previous studies (Holland and Holland, 2006: Andrews et al., 2021) by constructing a detailed spatiotemporal map of cell cycle progression in the neural tube, which highlights a sudden polarisation of cell division to the anterior and posterior tips of the nascent nervous system at the mid-neurula stage, concomitant with elongation of the anteroposterior axis. This includes discrete mitotic domains in the presumptive cerebral vesicle, and the chordoneural hinge of the tailbud, extending anteriorly into the posterior floor plate. By marking and following these cells in pulse-chase experiments, we demonstrate that anterior proliferative cells locally increase cell number within the cerebral vesicle in a ventral-to-dorsal temporal sequence, and that this wave of cell division is a pre-requisite for the emergence of brain cell type diversity (Figure 7A). In contrast, we show that proliferative cells in the chordoneural hinge and posterior neural plate generate floor plate progenitors that extend across 1/3rd of the anteroposterior axis. We further show that these posterior progenitors can be traced back to the dorsal blastopore lip of the gastrula, where they lie adjacent to posterior somite progenitors (Figure 7B). Finally, we find that the polarized cell division dynamic collapses prior to the onset of larval life, when proliferation resumes throughout the nervous system.

**Figure 7.**
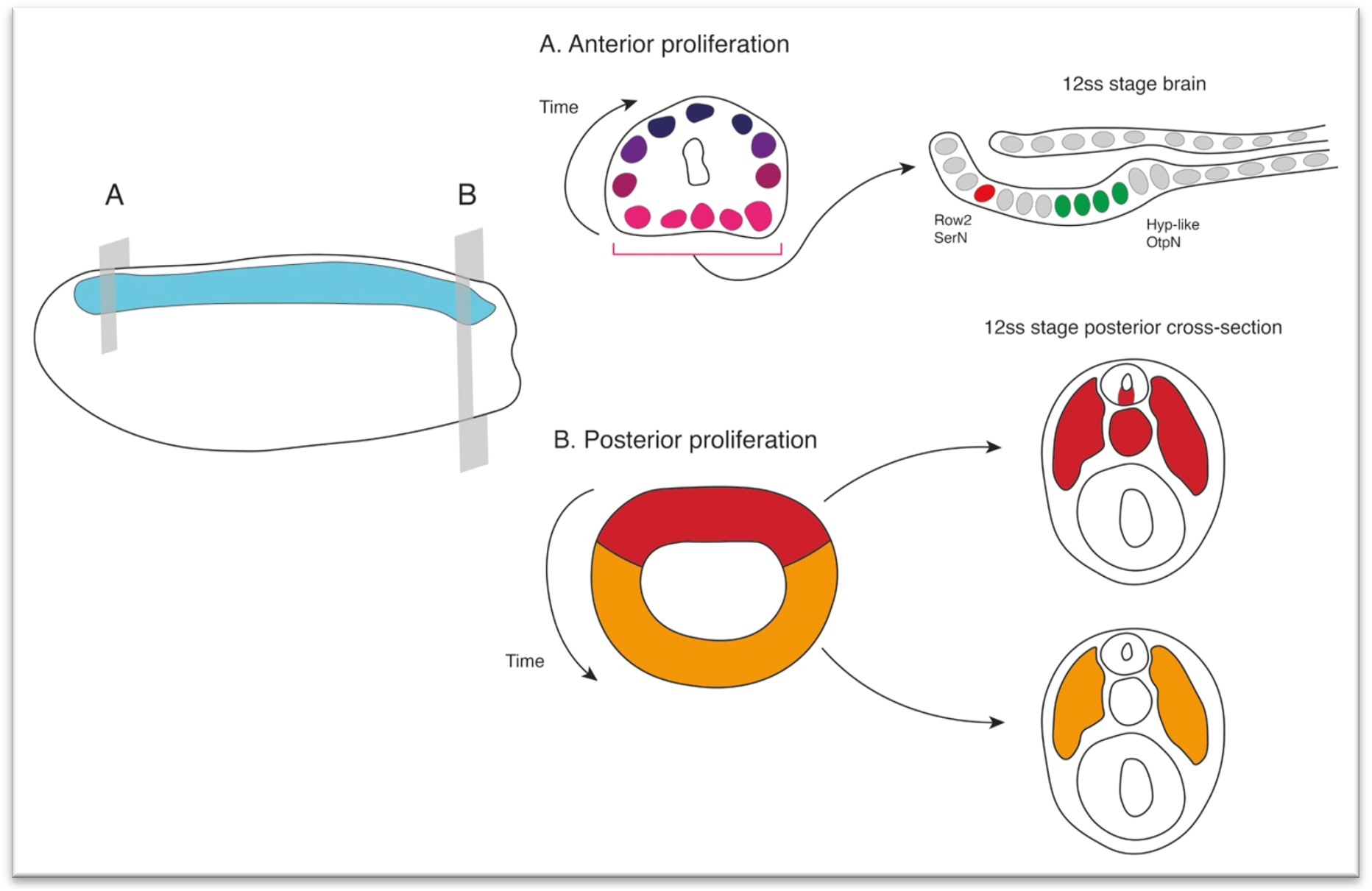
Contributions of cell proliferation to nervous system and axial development. During neurulation, cell division is restricted to the anterior and posterior poles of the neural plate. **(A)** Anteriorly a ventral to dorsal proliferative gradient in the cerebral vesicle is required to specify Row2 serotonergic neurons of the frontal eye and hypothalamic-like *Otp*-positive neurons. **(B)** Posteriorly, early proliferating cells in the dorsal blastoporal region contribute to the floor plate, notochord and somites, while progenitors in the ventroal and lateral regions of the blastopore divide later and are incorporated into the posterior somites.

The pattern of cell division we describe in amphioxus differs from what is seen in the vertebrate CNS in that the neuroectoderm of vertebrates proliferates extensively throughout its development. Early neural progenitors (neuroepithelial and radial glial cells) are known to undergo symmetrical divisions across the anteroposterior length of the neural plate and early neural tube (Qian et al., 2000; Götz and Huttner, 2005; Yingling et al., 2008). These divisions are used to expand the repertoire of stem cells that will later give rise to neurons and glia (Qian et al., 2000; Noctor et al., 2004). In fact, the rate of proliferation of these cells directly affects the size and regionalization of adult mammalian brains. For example, disruption of early signalling (e.g. FGF and Wnt/ Bcatenin) controlling neural tube proliferation in mammals deeply affects the size, thickness and convolution of the brain (Vaccarino et al., 1999; Chenn and Walsh, 2002; Megason and McMahon, 2002). In contrast, in other invertebrate chordates like ascidians, cells have been observed proliferating in the larval sensory vesicle and, to a lesser extent, in the visceral ganglion, thereby resembling more closely the anterior proliferation we observed in amphioxus. However, proliferation is absent posteriorly in the ascidian neural tube, indicating that neural morphology in these animals is primarily driven by cell re-arrangement and cell shape change (Tarallo and Sordino, 2004; Nakayama et al., 2005). This diversity of proliferative strategies in the different chordate subphyla supports the idea that tweaks in the timing and magnitude of cell division have played important roles in the emergence of neural complexity during chordate evolution.

Interestingly, we observed that proliferation in the trunk neural tube at 14 ss occurs specifically in cells of the floor plate and medio-lateral cells. Both cell types have recently been described as having a glial-like molecular signature in the early larva (Bozzo et al., 2021). Bozzo and collaborators speculate that proliferation of medio-lateral cells expressing the glial marker EAAT2 results in the formation of ependymal and ependymoglial cells in late larva stages (Lacalli and Kelly, 2002). Our results here provide experimental support for this hypothesis: while ventro-lateral cells already express neural markers, such as neurotransmitter synthesis and transport proteins (Candiani et al., 2012) and member of the ADAR family with function in RNA-editing (Zawisza-álvarez et al., 2020), medio-lateral cells are still dividing at the beginning of the larval stage and could therefore represent progenitors that maintain a glial signature and have a supporting role in the larval neural tube. Supporting this idea, we find that the transcription factor *Otp* is expressed in ventro-lateral cells of the trunk neural tube that are not proliferating at the early larva stage.

### Localised proliferation in the amphioxus brain is key for the proper formation of the frontal eye and hypothalamus

Anterior proliferative cells remain confined to the amphioxus cerebral vesicle, but cell division occurs sequentially in a ventral-to-dorsal direction, thereby contributing to different areas of the brain at distinct time points in development. This pattern suggested that the proliferation dynamics might have a role in regionalising the amphioxus brain, by contributing cells at specific times fated to particular types. In order to test such hypothesis, we pharmacologically arrested proliferation, at the time when this early regionalisation of the brain starts, coinciding with the time at which proliferation resumes and restricts to the cerebral vesicle (and chordoneural hinge posteriorly). We found that localised proliferation in the amphioxus brain is necessary to generate the anterior serotonergic cells of the frontal eye complex, as we previously proposed (Benito-Gutiérrez et al., 2018). In stark contrast, we found that glutamatergic neurons, including those anterior cells associated to the frontal eye complex, remain in place despite the absence of proliferation. This indicates that glutamatergic cells are likely to be born very early in development, but they only differentiate at later stages. Altogether, our results suggest that proliferation plays a key role in the formation of the frontal eye complex in amphioxus, by being essential to provide the right cell type architecture.

We also found that proliferation is indispensable to generate anterior brain *Otp* cells in amphioxus. *Otp* cells elsewhere in the neural tube were unaffected when proliferation was arrested, indicating that these cells were born earlier in development and differentiated later. This is congruent with the expression of *Otp* at 6-7ss, which showed clusters of *Otp* cells within the neural tube, at a stage immediately preceding our proliferation arrest assays. Apart from these neural tube clusters, no expression of *Otp* has been previously published elsewhere in the amphioxus nervous system. Here we report that *Otp* is specifically expressed in the amphioxus brain from the 10 ss onwards: first in a pair of cells posteriorly located within the posterior *Six3/6* domain; and later, at 12 ss, in two groups of 7-8 ventrolateral cells within the intercalated *Six3/6*-negative region. By combining *Otp* detection, EdU pulse-chase and cell-cycle arrest experiments we were able to birth-date all of these brain *Otp* cells. We found that the posterior pair of *Otp* neurons are indeed the first ones to be born, as they remain unaffected by the HU treatment given to embryos at the 6ss. Consistently with this, no EdU incorporation was detected in these *Otp* positive cells between the 6ss and 12ss, meaning they were born prior to the 6ss and differentiated at the 10ss. Strikingly, we did not detect Edu in the anterior *Otp* cluster at the 12 ss either, when it first appears. Given that this entire anterior domain disappears when proliferation is arrested, we reasoned that these cells must be born earlier in development, but not earlier than the posterior pair of Otp cells. Using an Edu pulse and chase strategy we found that these brain *Otp* progenitors are born between the 7 ss and the 8 ss stage but only differentiate at 12 ss.

Finally, at 12ss proliferation resumes in the posterior *Six3/6*-positive domain, in cells adjacent to the posterior *Otp* pair; these dividing cells likely contribute to the increase of posterior brain *Otp*-positive cells that are visible at 14ss, as these cells disappeared following inhibition of proliferation to this stage.

Interestingly, *Otp* is key to the development of the hypothalamic neuroendocrine system in all vertebrates (Wang and Lufkin, 2000; Del Giacco et al., 2008). As in amphioxus, this area of the brain is located in vertebrates posteriorly to the preoptic area, in amphioxus defined by the frontal eye complex, here shown through *SerT* expression (Fig4). In ascidians, *Otp* cells have been shown to localise in a similar anterior position within the sensory vesicle, also adjacent to surrounding Six3/6-positive cells (Moret et al., 2005). Altogether, our results strongly suggest that brain *Otp* cells might be revealing a putative hypothalamic region within the amphioxus brain that develops through proliferation and growth from the intercalated Six3/6-negative region around the 7-8ss.

### The amphioxus tailbud is a source of posterior floor plate and notochord cells

In the posterior body, we found using EdU pulse-chase analysis that proliferative cells in the chordoneural hinge of the tailbud specifically contribute to the elongation of the floor plate and notochord. This is mediated by a burst of cell division occurring in the mid-neurula after 6 ss, when proliferation restricts to the most posterior and anterior ends of the neural tube. Prior to this developmental stage, cells proliferate in the posterior neural plate throughout axial development, and pass sequentially through S-phase and mitosis. This means that the posterior neural plate is an active proliferative domain from gastrulation to the end of somitogenesis. Given this, we were able to track the progeny of dividing axial progenitors in the blastopore via a series of EdU pulse-chase experiments. With this, we demonstrate that midline axial progenitors originate from the dorsal blastopore lip of the late gastrula. There, they lie adjacent to progenitors of the posterior somites, which are located in the lateral and ventral blastopore lips.

As such, the location and fates of these axial progenitor cells in amphioxus bears strong similarity to the dynamics of midline progenitor cells identified in vertebrates. In amniotes, homotopic grafting approaches and direct cell marking have located midline progenitors of the floor plate and notochord that arise in the node-streak border of the gastrula, and undergo clonal expansion within the chordoneural hinge (Catala et al., 1996; Cambray and Wilson, 2002; Mugele et al., 2018). Similarly, fate mapping of the dorsal blastopore lip in *Xenopus* has located derivatives in the posterior notochord in floor plate (Gont et al., 1993). In all cases, midline progenitor cells further extend the tissue primordia formed during gastrulation and do so to a degree that is species-specific, aligning with variations in nutritional supply (O’Farrell, 2015; Steventon et al., 2016). In externally developing anamniotes, axial progenitors undergo little clonal expansion and make minor contributions to axial length, whereas in amniotes they undergo extensive clonal expansion and growth, thereby driving a significant posterior enlargement of the embryo (Steventon et al., 2016; Attardi et al., 2019). Our results in amphioxus therefore support a model in which midline axial progenitors may be an ancestral trait in chordate embryogenesis, in which evolutionary transitions in clonal dynamics have facilitated divergent contributions to axial length.

## Conclusions

Our work provides a spatiotemporal map of cell cycle progression during the early development of amphioxus (up to the early larval stage). Proliferation is widespread in the archenteron throughout the whole period observed. However, by focusing our analysis on cell division within the CNS, we find that the nervous system is spatially segregated in three different developmental modules characterised by distinct cell division dynamics. Anteriorly, within the brain, proliferation acts locally to increase the neural type repertoire at specific time points. We find that cell proliferation is necessary to fully equip the frontal eye with the correct neurotransmitting cells, before the eye spot fully develops into a functional photoreceptor able to integrate sensory information (Vopalensky et al., 2012). In the brain, proliferation is also needed to grow the ventral portion of the brain, which we identify here as hypothalamic based on *Otp* expression. It is tempting to speculate that this phase of hypothalamic development is preparatory for entering the subsequent feeding phase, when a neuroendocrine system would need to be in place. In parallel, neuronal cells gradually differentiate in the trunk nervous system, independently of cell division, and posteriorly, tailbud progenitors contribute to elongating the posterior neural axis, which at the period analysed remains generally undifferentiated. Unexpectedly, the amphioxus body plan is robust to cell cycle arrest. Embryos develop with fewer cells but they still harbour a dorsal neural tube with an underlying notochord and the correct number of somites, demonstrating that cell division is primarily required to confer a proper geometry and to increase neuronal type complexity.

## Author Contributions

Study design: GG, TGRA, EBG; acquisition and analysis of data: GG, TGRA; writing – original draft: GG, TGRA; manuscript review and editing: GG, TGRA, EBG. Project administration: EBG. All authors have read and agreed to the published version of the manuscript.

## Acknowledgments

The authors would like to thank Matt Wayland from the CAIC for assistance during imaging; and to Hector Escrivà and Jordi Garcia-Fernàndez for helping with sourcing amphioxus during the pandemic difficult times.

