## Supplementary Information for "Restricted proliferation during neurogenesis contributes to regionalization of the amphioxus nervous system"

### Supplementary Figure1

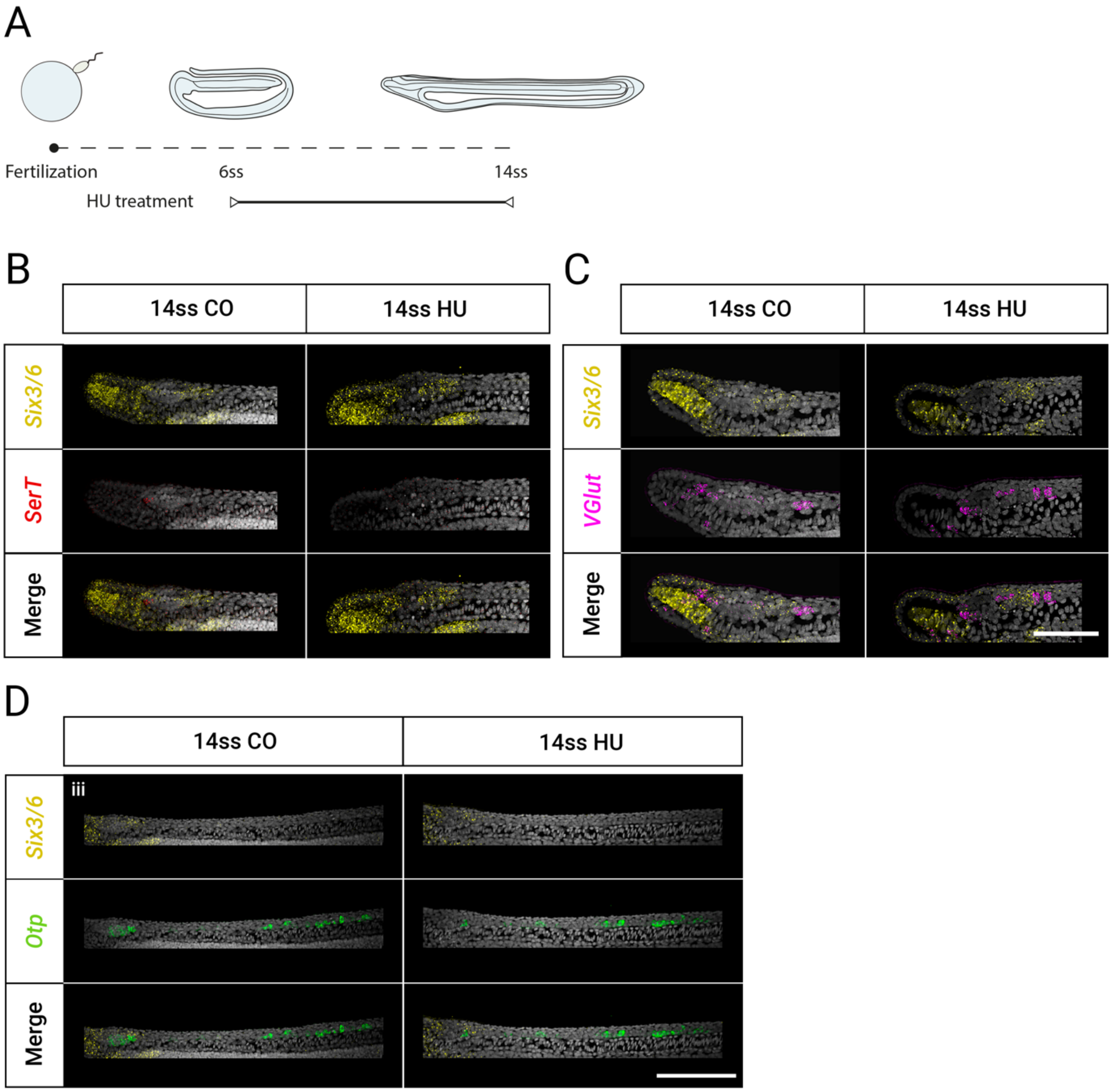


**Supplementary Figure 1. Gene expression following inhibition of proliferation to the 14ss stage.** (A) Experimental design of late hydroxyurea treatments used in this study. (B) Expression of *Six3/6* (yellow) *SerT* (red) and *VGlut* (magenta) remains similar to 12ss and responds in the same way to inhibition of proliferation: *SerT* expression in the cerebral vesicle is lost while *Six3/6* and *VGlut*-positive cells persist in HU-treated embryos. (D) *Otp (green)* expression is lost in the anterior clusters in the *Six3/6*-negative domain, and is reduced to only a pair of cells in the posterior *Six3/6*-positive domain following HU treatment. Scale bar: 50µm (B-C); 100µm (D).
